# Characterizing key osmolytes and osmoprotectants in drought-stressed Scotch pine: a differential approach

**DOI:** 10.64898/2026.03.23.713677

**Authors:** Alexander V. Kartashov, Ilya E. Zlobin, Yury V. Ivanov, Alexandra I. Ivanova, Anastasia Orlova, Nadezhda Frolova, Alena Soboleva, Svetlana Silinskaya, Tatiana Bilova, Andrej Frolov, Vladimir V. Kuznetsov

**Author notes:** Corresponding author: Alexander V. Kartashov, K.A. Timiryazev Institute of Plant Physiology Russian Academy of Science, 35 Botanicheskaya st., Moscow, 127276, Russia.

## Abstract

During drought, numerous compounds accumulate in plant tissues, but their physiological roles remain unclear – they may function as osmolytes, osmoprotectants, or merely arise as by-products of stress-induced metabolic shifts. We developed an experimental approach to link accumulation patterns with specific functions, using Scots pine (*Pinus sylvestris* L.) saplings subjected to water deprivation and subsequent rewatering as a model system. We monitored changes in relative water content (RWC) and osmotic adjustment dynamics, employed untargeted primary metabolite profiling for preliminary screening of compounds correlated with water status, and performed quantitative GC-MS and LC-MS analyses of selected metabolites. Major inorganic cations (K⁺, Ca²⁺, Mg²⁺) were also quantified to assess their potential roles. Our results revealed that tryptophan, valine, and lysine – though generally present in low abundance – exhibited selective accumulation under severely reduced RWC (≤ 70%), suggesting their involvement as osmoprotectants. Major organic acids, particularly shikimic acid, showed trends consistent with osmotic adjustment. Notably, neither sucrose nor inorganic cations appeared to function as primary osmolytes in this context. The proposed approach offers a viable strategy for identifying compounds involved in plant adaptation to water deficit, with potential applications in breeding programs aimed at improving drought tolerance.

**Highlights:** An approach to identify osmolytes and osmoprotectants was implemented Accumulation of Trp, Val and Lys was consistent with their role in osmoprotection Osmotic adjustment relied predominantly on organic acids than on inorganic ions Monosaccharides but not sucrose correlates with changes in needle water status

## 1. Introduction

Water deficit poses a critical threat to the majority of terrestrial plant biomes, and its significance is escalating in the context of rising global temperatures, thereby underscoring the urgency of developing more drought-resistant tree varieties (Polle *et al*., 2019). Under drought conditions, the primary determinant of tree survival is the capacity to maintain tissue hydration and prevent desiccation-induced mortality (Anderegg *et al*., 2016; Choat *et al*., 2018). Trees can enhance hydraulic safety through a suite of structural adaptations, including modifications in xylem embolism tolerance, hydraulic conductivity, and the leaf-to-sapwood area ratio (reviewed in (Rowland *et al*., 2023)). Nevertheless, these structural traits are inherently "hard" or growth-based, limiting their capacity for rapid and reversible adjustment in response to fluctuating water availability. A defining feature of drought stress is its unpredictability: droughts may vary in timing, duration, and intensity, occur as isolated or consecutive events within a single season, and, in the case of perennial trees spanning multiple seasons, exhibit compounded temporal variability. Consequently, breeding strategies aimed at enhancing drought tolerance in trees may benefit from a focus on "soft" traits – those capable of dynamic and reversible modulation in accordance with changes in water supply. Such an approach holds promise for accelerating the development of climate-resilient tree varieties and ensuring the long-term sustainability of forestry in an increasingly water-limited future.

Among the most important “soft” mechanisms of drought adaptation is osmotic adjustment – the active accumulation of solutes in cells in response to water deficit (Osmolovskaya *et al*., 2018; Turner, 2018) – as opposed to the passive increase in solute concentration resulting from cellular water loss. Osmotic adjustment represents a universal and predominant mechanism for maintaining turgor under water-limited conditions, surpassing the contributions of modifications in apoplastic water content or cell wall rigidity (Bartlett *et al*., 2012). The accumulation of soluble carbohydrates (though not total non-structural carbohydrates) in tree branches and stems is positively influenced by seasonal aridity worldwide (Blumstein *et al*., 2023). In leaves of drought-stressed plants, concentrations of soluble sugars are maintained despite substantial reductions in starch content, as demonstrated in a meta-analysis by (Wang and Wang, 2023). These findings align with the hypothesis that maintaining an appropriate osmotic potential is of central importance for plants experiencing water deficit. Indeed, insufficient capacity for osmotic regulation has been identified as a likely contributor to tree mortality in models simulating tree growth and metabolism (Potkay *et al*., 2022). The process of osmotic adjustment itself appears to depend more on the overall capacity to lower osmotic potential than on the specific composition of osmotically active compounds (Zivcak *et al*., 2016). Nevertheless, different plant species employ a remarkably broad spectrum of osmolytes potentially involved in osmotic adjustment (Slama *et al*., 2015), and the factors governing the selection of particular compounds by a given species remain poorly understood. Compounding this complexity, the pronounced accumulation of a given metabolite under water stress does not necessarily imply its direct involvement in osmotic adjustment. Drought disrupts multiple physiological processes to varying degrees (Singh *et al*., 2020), and such heterogeneous disturbances alter the balance between synthesis, turnover, and consumption of various metabolites. Consequently, shifts in metabolite concentrations under stress may reflect passive metabolic imbalances rather than actively regulated osmotic adjustment.

Under severe water deficit, stress can progress to a degree that compromises the proper hydration of macromolecules and cellular structures (Zivcak *et al*., 2016). Leaf relative water content (RWC) serves as a direct indicator of cellular hydration status (Martinez[:Vilalta *et al*., 2019), with values in the range of 70–75% likely representing a critical threshold. Below this level, plants typically experience loss of turgor, disruption of biochemical processes, and diminished capacity for tissue rehydration upon rewetting (Lawlor and Cornic, 2002; Bartlett *et al*., 2012; John *et al*., 2018; Trueba *et al*., 2019). Therefore, when RWC declines below these thresholds, plants are expected to activate protective mechanisms aimed at preserving cellular integrity, primarily through the accumulation of stress-protective compounds, or osmoprotectants. Osmoprotectants are low-molecular-weight, highly soluble, and electrically neutral molecules of diverse chemical nature (Zivcak *et al*., 2016), capable of stabilizing macromolecular structures and mitigating stress-induced functional impairments (Sharma *et al*., 2021). Major classes of osmoprotective compounds include methylamines, polyols (including sugars such as sucrose and trehalose), as well as specific amino acids and their derivatives. These compounds share the abilities to promote protein folding and protein solubility to a different extent (Rabbani and Choi, 2018). For instance, the accumulation of proline is frequently observed in water-stressed plants and is thought to serve primarily a protective role in stabilizing cellular structures rather than contributing substantially to osmotic adjustment (Blum, 2017). However, while the osmoprotective effects of numerous compounds have been unequivocally demonstrated *in vitro* (see e.g. (Sharma *et al*., 2021)), establishing their functional relevance *in vivo* remains considerably more challenging. Consequently, the adaptive rationale underlying the diversity of osmoprotective compounds employed by different organisms – a question paralleling the earlier discussion of osmolyte diversity – remains largely unresolved (Rabbani and Choi, 2018).

The rapid advancement of metabolomics has enabled the simultaneous analysis of concentration changes in tens to hundreds of compounds in water-stressed plants (Kumar *et al*., 2021). However, a fundamental question remains: how can one determine whether an increase in a given compound under water deficit reflects its active involvement in osmotic adjustment or osmoprotection, or merely represents a passive consequence of metabolic disturbance? If the objective is to breed tree varieties with enhanced capacity to maintain osmotic balance under limited water availability, it is essential to specifically target compounds that participate in osmotic adjustment. Achieving this requires the ability to distinguish between metabolites involved in osmotic adjustment, those contributing to stress protection, and those whose concentrations shift as incidental by-products of stress-induced metabolic alterations (Zulfiqar *et al*., 2020). We argue that rather than focusing solely on differences in mean concentrations of osmotically active compounds under drought, it is more informative to examine the relationships between compound concentrations and plant water status parameters at the individual plant level. Based on this rationale, we propose the following framework linking the physiological functions of compounds to their accumulation patterns (see also (Zivcak *et al*., 2016)):

- **Osmoregulators** would exhibit correlations with changes in plant water and osmotic status, with such relationships being more pronounced under water-stressed than under non-stressed conditions. These compounds are expected to be present at relatively high concentrations, given that osmoregulation is a "bulk" property dependent on the total solute concentration in cellular compartments.;
- **Osmoprotectants** would show a distinct pattern of accumulation when leaf relative water content (RWC) falls below 70–75%, a threshold associated with substantial dehydration of the intracellular milieu. These compounds may accumulate to relatively low concentrations compared to osmoregulators (Blum, 2017);
- the remaining compounds involved in neither osmotic regulation nor stress protection would not correlate with changes in osmotic status or RWC and are thus unlikely to be of direct relevance for breeding efforts.

A species with high phenotypic plasticity and a broad ecological range would serve as an optimal model for testing these proposed relationships. Therefore, Scots pine (*Pinus sylvestris* L.) was chosen as a model due to its well-documented high potential for physiological adaptation to water deficit. Its choice is further justified by its substantial economic importance and its status as a dominant forest-forming species across the European part of Russia and Northern Europe. Guided by the functional classification outlined above, we aimed to identify compounds specifically involved in osmoregulation and osmoprotection in needles of water-stressed *P. sylvestris* L. saplings, in which pronounced drought effects on water status had previously been characterized (Zlobin *et al*., 2022). The identification strategy comprised three main steps. First, the dynamics of osmotic adjustment in needles were characterized. Second, untargeted metabolite profiling was performed to identify correlations between changes in relative metabolite abundances and parameters of plant water status. Third, targeted quantitative analysis of metabolites identified in the previous step was conducted, alongside measurements of major inorganic cations (K^+^, Ca^2+^, Mg^2+^), to evaluate their potential roles as osmolytes or osmoprotectants.

## 2. Materials and methods

### 2.1. Chemicals

Materials were obtained from the following manufacturers: Ecos-1 (Moscow, Russia): hexane (puriss p. a.); Macherey-Nagel GmbH and Co KG (Düren, Germany): *N*-methyl-*N*-(trimethylsilyl)trifluoroacetamide (MSTFA, MS grade); Vekton (Saint-Petersburg, Russia): methanol (LC grade). All other chemicals were purchased from Merck KGaA (Darmstadt, Germany).

### 2.2. Plant material and experimental conditions

Plant material and experimental design were previously described in (Zlobin *et al*., 2022) and (Zlobin *et al*., 2023). Briefly, containerized three-year-old saplings of Scots pine (*Pinus sylvestris* L.) grown from improved seeds collected in the Semyonovskoe Division of Forestry (Nizhny Novgorod Region, Russia) were obtained from the forest nursery “Semenovskij specsemleskhoz” (Nizhny Novgorod Region, Russia). Saplings were transplanted to 7 L-pots filled with soil substrate (peat, sand, ground limestone, 350 mg/L, N, 30 mg/L P, 400 mg/L K, pH 5.5–6.0). Following an overwintering period outdoors, the plants were acclimated in a cold frame at approximately 5°C for two weeks before being transferred to a heated greenhouse under natural illumination for the duration of the experiment.

The plants were divided into two groups: the first group was watered twice weekly to full soil capacity (“control” variant), while the second group was subjected to water deprivation (“drought” variant) (Supplementary information, Figure S1).

**Fugure S1.**
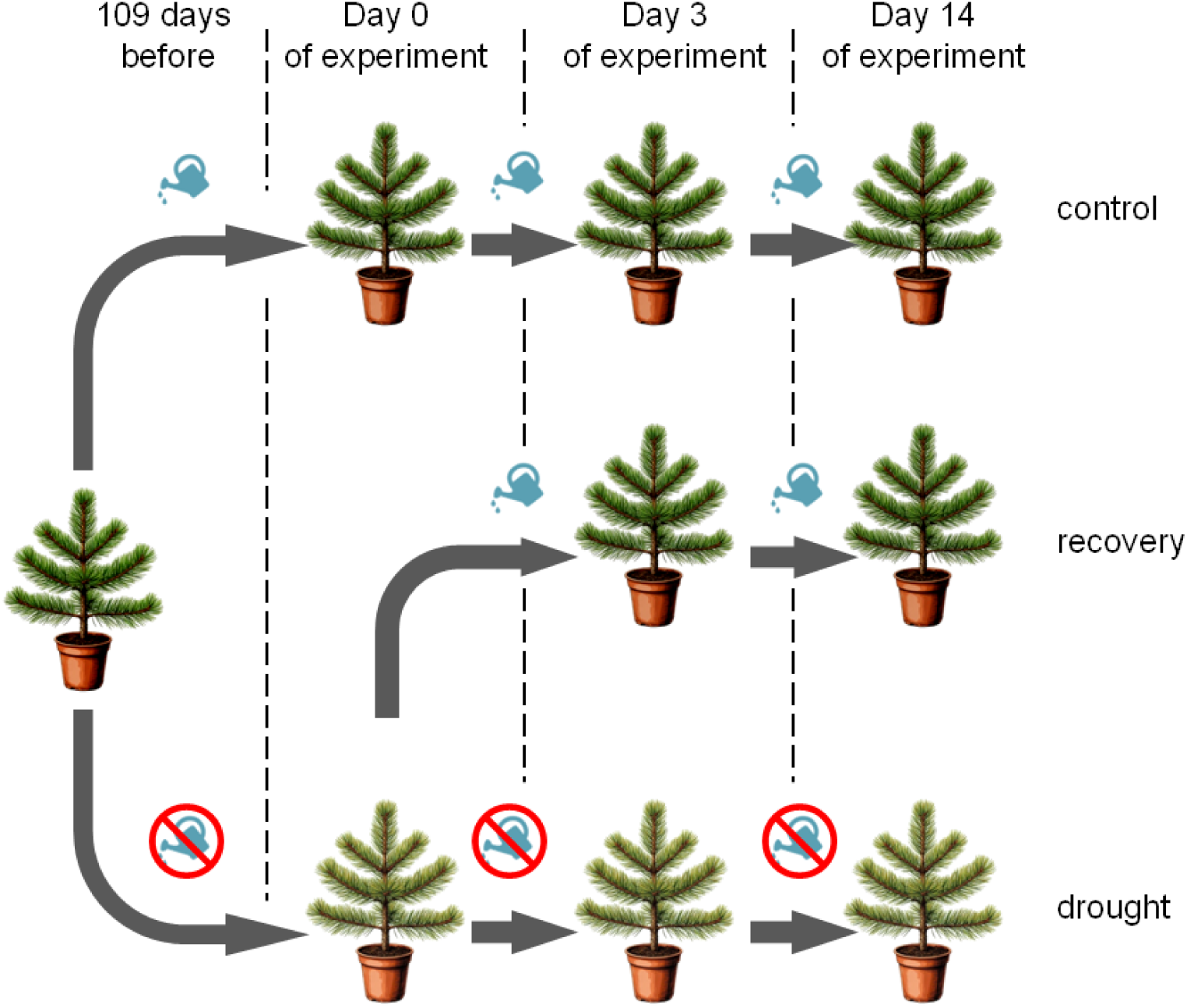
Experimental design used to impose drought stress and subsequent rewatering Scots pine saplings.

After 109 days of water deprivation, the latter group exhibited substantial water stress (see (Zlobin *et al*., 2022)), this time point was the designated as Day 0 of experiment. On Day 0, relative water content (RWC) and absolute water content (WC) and osmotic potential were determined in upper and lower needles needles both control and drought-stressed plants, with six biological replicates per each variant. Osmotic potential was measured in cell sap squeezed from needles on ice using a cryo-osmometer Osmomat 030 (Gonotec, Germany). Subsequently, half of the drought-stressed plants were rewatered to full soil capacity (“recovery” variant), while the remaining half subjected continuous drought. After three (Day 3) and fourteen (Day 14) days, RWC, WC, and osmotic potential were determined in upper and lower needles of control, drought-stressed and recovering plants (six biological replicates per treatment per time point). Additionally, six biological replicates of upper and lower needles from each experimental variant at each time point (Day 0, Day 3, Day 14) were collected, flash-frozen in liquid nitrogen and stored at -70°C for metabolomic analysis and determination of inorganic ion content.

### 2.3. Calculation of osmotic adjustment

To quantify osmotic adjustment, it was necessary to distinguish between the change in osmotic potential attributable to solute concentration resulting from water loss and that attributable to active solute accumulation (Turner, 2018). Two methods of correction for changes in water content were employed.

**In Method 1**, osmotic adjustment (OA1) was calculated according to (Babu *et al*., 1999) with modifications. Changes in RWC in each plant, relative to the mean RWC of control plants at the corresponding time point (Day 0, 3, or 14), were used to correct osmotic potential for water loss:

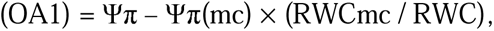

where OA1 – calculated osmotic adjustment; Ψπ – osmotic potential in a given needle sample; Ψπ(mc) – mean osmotic potential in control plants at a given time point (0 days, 3 days, 14 days); RWCmc – mean RWC in control plants at a given time point (0 days, 3 days, 14 days); RWC – relative water content in a given needle sample.

**In Method 2**, osmotic adjustment (OA2) was calculated according to (Alves and Setter, 2004) with modifications. Changes in absolute water content in each plant’s needles, relative to the mean absolute water content of control plants at the corresponding time point, were used for correction:

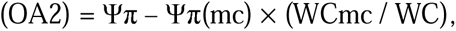

where OA2 – calculated osmotic adjustment; Ψπ – osmotic potential in a given needle sample; Ψπ(mc) – mean osmotic potential in control plants at a given time point (0 days, 3 days, 14 days); WCmc – mean absolute water content in control plants at the corresponding time point; WC – absolute water content in a given needle sample.

### 2.4. Untargeted profiling of primary metabolites

Primary thermostable and thermolabile metabolites were analysed by gas chromatography coupled on-line to quadrupole mass spectrometry with electron ionization (GC-EI-Q-MS) and ion-pair reversed phase high-performance liquid chromatography coupled online to triple quadrupole tandem mass spectrometry (RP-IP-HPLC-QqQ-MS/MS) in multiple reaction monitoring (MRM) mode, respectively, with all sample preparation and extraction procedures accomplished as described by (Shumilina *et al*., 2023). The specific instrumental settings employed for corresponding GC-MS and LC-MS experiments are listed in Supplementary information 2, (Tables S2.1 and S2.2, respectively).

Processing of the GC-MS data and relative quantification of individual analyte abundances relied on Automated Mass Spectral Deconvolution and Identification System, AMDIS v.2.66 and MSDial. Identification of trimethylsilyl (TMS) and methyl oxime (MEOX)-TMS derivatives of the analytes relied on co-elution with authentic standards or/and spectral similarity search against available spectral libraries – NIST (National Institute of Standards and Technology, https://webbook.nist.gov/chemistry/), GDM (Golm Metabolome Database, http://gmd.mpimp-golm.mpg.de/) and in-house spectral library ASL (Authentic Standard Library). Annotation of individual analytes in LC-MS chromatograms relied on co-elution with authentic standards in corresponding MRM scans, whereas relative quantitative analysis assumed integration of corresponding chromatographic peaks at specified transitions and tRs – with PeakView 2.2 and MuliQuantTM 3.0.2. softwares from AB Sciex (Darmstadt, Germany). After processing, the data matrices (including peak areas at specified tRs) obtained from the LC-MS and GC-MS analyses were combined and post-processed.

Prior to the statistical analysis, the data were normalised to the dry mass of samples (DW normalisation). Data post-processing relied on the methods of multivariate, and invariant statistics implemented in the Metaboanalyst 6.0 on-line tool (https://www.metaboanalyst.ca/). A detailed description of data post-processing is provided in Supplementary information 2. The results of the analysis were corrected for false discovery rate (FDR) using the Benjamin-Hochberg method at *p* ≤ 0.05 and fold change (FC) threshold of ≥ 2.

### 2.5. Absolute quantification of selected primary metabolites

Based on the results of the initial metabolite profiling, a set of individual metabolites with characteristic drought and/or post-drought recovery-dependent behaviour (in total 10 thermally stable metabolites) was addressed in more detail. Targeted absolute quantitative analysis relied on a standard addition method for low-abundant metabolites and an external calibration method for highly abundant metabolites. For these ten authentic standards (fumaric acid, malic acid, shikimic acid, ascorbic acid, fructose, glucose, sucrose, lysine, valine, tryptophan) prepared as a total mix serially diluted in the range from 0.5 pmol/μL to 0.1 nmol/μL, were used. To assess the contents of individual metabolites in the most reliable way, both analytical methods were employed.

The quantitative data were expressed in mol per g DW as mean ± standard error (SE) of six biological replicates. All calculations were done within the linear dynamic ranges (LDRs, *R*^2^ ≥ 0.95) derived in additional dilution experiments (n=3) with individual authentic standards. A detailed description of data processing provided in Supplementary information 2.

To assess whether metabolite concentrations changed significantly under the experimental conditions, Z-scores were calculated using a critical value of 1.96. This approach was adopted because the marked differences in metabolite concentrations among control plants at Day 0, Day 3, and Day 14 prevented direct comparison of the experimental groups with their respective time-point controls.

### 2.6. Major inorganic cations contents analysis

The contentsof K^+^, Ca^2+^, Mg^2+^ were determined by flame atomic absorption spectrometry (AAS) (Ivanov *et al*., 2024). Fresh needles probes were dried at 80 °C for three days until a constant weight was achieved. Approximately 50 mg of dry sample was digested in a mixture of concentrated HNO_3_ and HClO_4_ (2:1 v/v). Subsequently, 37 % H_2_O_2_ was added to the cooled samples until clarification and cessation of foaming. The metal contents were measured using an atomic absorption spectrophotometer (AA-7000, Shimadzu, Japan) equipped with a hollow-cathode lamps (Hamamatsu, Japan). Prior to the analysis, samples were appropriately diluted with 0.1M HNO_3_. For the measurement of Ca and K, the diluting solutions included LaCl_3_ (10 g La^3+^ per L) or LiCl (1 g Li^+^ per L) respectively to mitigate ionization interference. Parallel sample blank probes were prepared in same manner for background correction.

### 2.7. Statistical analysis

Each sapling was treated as a biological replicate; accordingly, there were 6 biological replicates for each experimental variant at each time point. Statistical processing of metabolomiс data is given in corresponding sections. For plant physiology parameters (RWC, OA1, OA2) and major inorganic cations contents data were statistically analyzed using SigmaPlot 12.3 (Systat Software, USA). Pearson’s correlation coefficients were calculated to assess the strength and significance of relationships between variables, and those presented in the text are significant at *p* ≤ 0.05. Values presented in figures are arithmetic means, and values presented in tables are arithmetic means ± standard errors.

## 3. Results

### 3.1. Dynamics of relative water content and osmotic adjustment in pine needles

Parameters of needle water status were analyzed separately for needles sampled from upper and lower parts of the plants, as described in the Materials and Methods section. However, differences in water status between upper and lower needles were found to be minimal. At the onset of the two-week experimental period (Day 0), drought-treated plants had already developed a significant water deficit. In lower needles of drought-stressed plants, RWC decreased to 75.9%, compared to 86.2% in control plants. In upper needles, RWC values were 77.6% and 87.8% in drought-stressed and control plants, respectively (Figure 1A). After two weeks of continued water deprivation (Day 14), RWC declined further to 68.6% in lower needles and 70.5% in upper needles of drought-stressed plants. In contrast, rewatering restored RWC to control levels by the end of the two-week experimental period (Figure 1A).

The extent and dynamics of osmotic adjustment differed between the two calculation methods employed. Osmotic adjustment calculated according to Method 1 (OA1) was 0.207 MPa in lower needles and 0.253 MPa in upper needles at the beginning of the drought period, increasing to 0.508 MPa and 0.442 MPa, respectively, by its conclusion (Figure 1B). Following rewatering, OA1 persisted at elevated levels, measuring 0.191 MPa in lower needles and 0.146 MPa in upper needles. In contrast, osmotic adjustment calculated according to Method 2 (OA2) yielded values more than twofold lower than OA1, reaching 0.150 MPa in lower needles and 0.206 MPa in upper needles of drought-stressed plants by the end of the two-week period (Figure 1B). Unlike OA1, rewatering resulted in the dissipation of OA2 to non-stress levels (0.021–0.027 MPa) by the end of the two-week recovery period.

It is plausible that OA1 values are more physiologically relevant than OA2, as prior exposure to water deficit can alter the absolute water content in tissues of well-watered plants through elastic and apoplastic adjustments (Bartlett *et al*., 2012). Consequently, correction based on RWC is likely advantageous compared to correction based on absolute water content. Nevertheless, both methods for calculating osmotic adjustment were employed concurrently in subsequent analyses.

**Figure 1.**
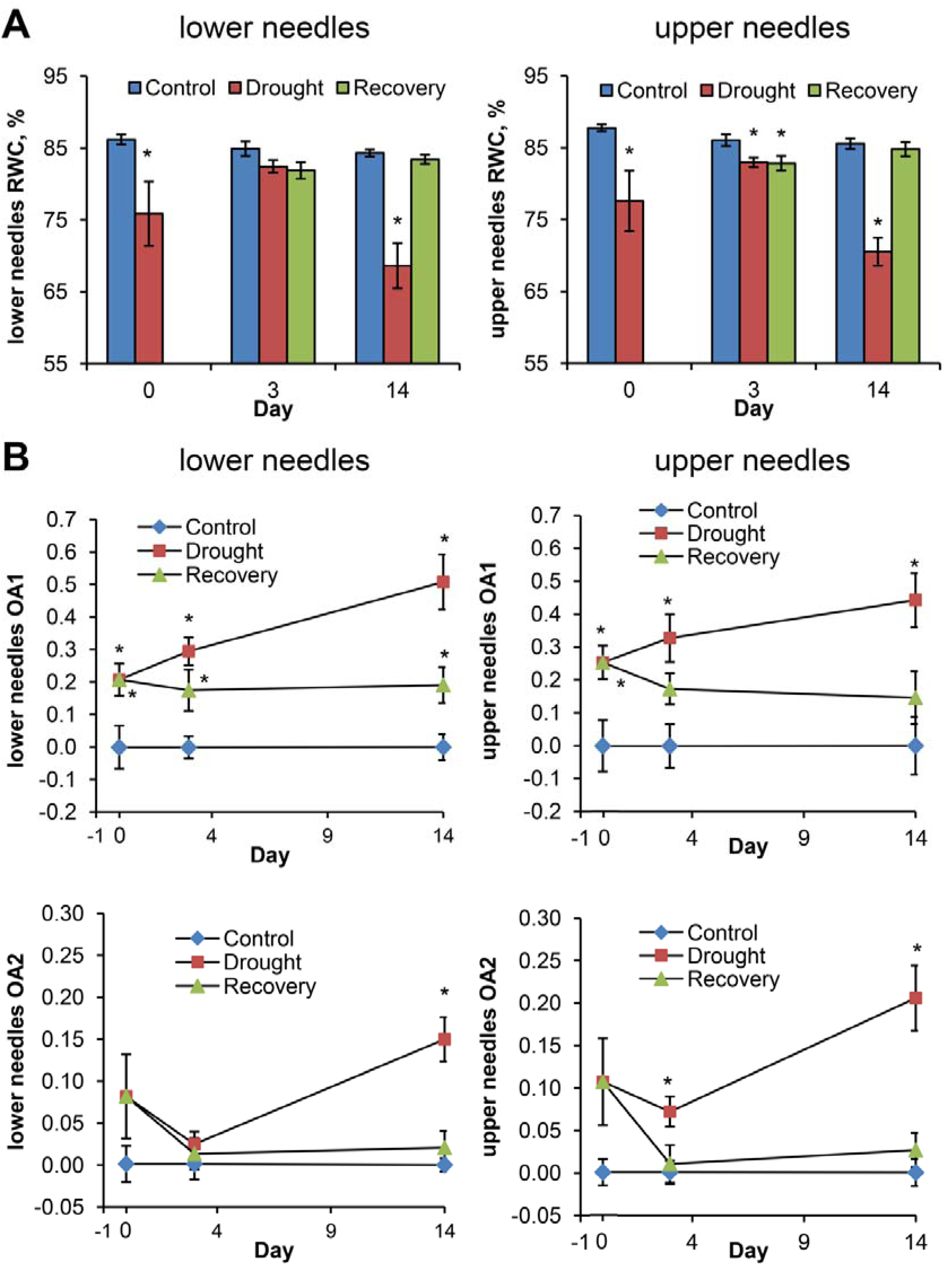
Dynamics of RWC (A) and osmotic adjustment (B) in upper and lower needles of *Pinus sylvestris* saplings. Pairwise comparisons were performed using Student’s *t*-test, asterisks (*) denote significant differences at p ≤ 0.05 from control and experimental variants at each time point Across the entire dataset, RWC correlated significantly with both OA1 (*r* = -0.597) and OA2 (*r* = -0.751) (Figure 2). In control plants, only OA2 exhibited a significant correlation with RWC (*r* = -0.501). In drought-stressed plants, both OA1 (*r* = -0.445) and OA2 (*r* = -0.727) demonstrated significant correlations with RWC. In recovering plants, no significant correlation between RWC and osmotic adjustment was detected, irrespective of the calculation method employed.

**Figure 2.**
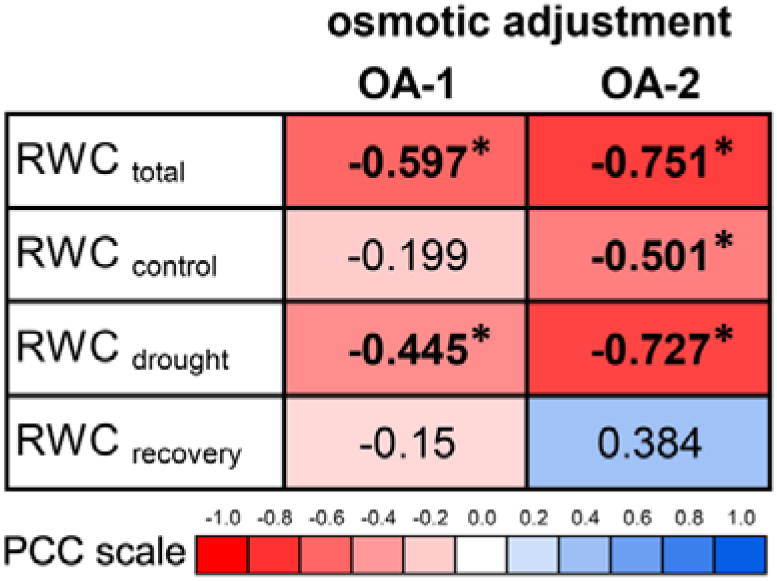
Correlations (Pearson’s correlation coefficients – *r*) between osmotic adjustment and RWC for total dataset, control, drought-stressed and recovering plants. Asterisks (*) denote significance of the Pearson’s correlation at *p* ≤ 0.05

### 3.2. Untargeted metabolomics analysis of the water deficit response

GC-EI-Q-MS-based profiling of aqueous methanolic extracts prepared from needles of drought-treated and control *P. sylvestris* saplings revealed a total of 122 individual polar, thermally stable metabolites that could be reliably annotated through spectral similarity searches and/or co-elution with authentic standards (Table S2.3). Among these, 86 individual compounds could be annotated to trimethylsilyl (TMS) and methoxyamine(MeOX)-TMS derivatives of specific polar metabolites, which mostly represented plant primary metabolism. As certain metabolites yielded multiple MeOX-TMS derivatives, the total number of identified (structurally annotated) metabolites appeared to be only 67. These metabolites represented 14 proteinogenic and non-proteinogenic amino acids, four amines, two phosphates, 13 organic acids, 24 sugars (including sugar phosphates and sugar alcohols), two polyphenolic compounds, two pyrimidines, two metabolites concerning to *myo*-inosytol and *myo*-inositol-phosphate, and four representatives of other classes. For an additional 36 analytes unambiguous annotation was not possible. Among them, 15 species could be annotated to specific metabolite classes (mostly sugars), while the remaining 22 features were classified as unknown metabolites and labelled accordingly with their unique retention indices (RIs) (Table S2.3).

The LC-MS/MS-based profiling of acidic aqueous *P. sylvestris* needles extracts relied on multiple reaction monitoring (MRM) experiments and annotation by co-elution with well-characterized set of authentic standards. This approach revealed in total 159 thermally labile primary metabolites annotated in the whole dataset (Table S2.4). These included 35 nucleosides and nucleotides/their derivatives, 25 amino acids, 36 sugars, sugar phosphates and their derivatives; 47 carboxylic acids; nine coenzyme A derivatives; three inorganic compounds, four representatives of other classes. After merging the GC-MS and LC-MS result sets, a combined matrix with 281 entries was built and processed with the MetaboAnalyst 6.0 online software tool.

To identify the metabolites responsive to alterations in water status, correlations between the changes in relative abundances of all individual metabolites present in the matrix and parameters of the needle water status (RWC, OA1, OA2) were calculated across the whole dataset (Figure 3; Table S3). For this analysis, data obtained from upper and lower needles across different experimental time points were combined and grouped into three categories: control (*n* up to 35), drought (*n* up to 36), and recovery (*n* up to 23). In addition, the total dataset (*n* up to 94) was analyzed. Metabolites were selected for further consideration if they exhibited a significant negative correlation with RWC and positive correlations with OA1 and/or OA2 in plants subjected to continuous drought and in the total experimental dataset. Among amino acids, the most prominent correlation with tissue water status was found for tryptophan, which demonstrated both negative correlation with RWC and positive correlation with levels of osmotic adjustment when analyzed by either GC-MS or LC-MS. Valine and lysine exhibited weaker but still significant correlations with water status in GC-MS analyses; however, these correlations were absent for lysine in LC-MS analyses, and valine was not detected by this method. Notably, proline – a well-established stress protector and marker of drought stress response – did not exhibit significant correlations with plant water status (Table S3). Among organic acids, both fumaric and malic acids showed correlations with the changes in plant water status when analyzed by GC-MS, but not by LC-MS. Sugars and sugar alcohols, in general, did not display any correlations with plant water status, with the exception of fructose quantified by GC-MS. Interestingly, two compounds with long aliphatic moieties - heptylaldehyde and farnesyldiphosphate - demonstrated relatively consistent correlations with water status (Table S3). However, given their hydrophobic nature, these compounds are unlikely to function as osmolytes or osmoprotectants in plants and were therefore not considered further.

**Figure 3.**
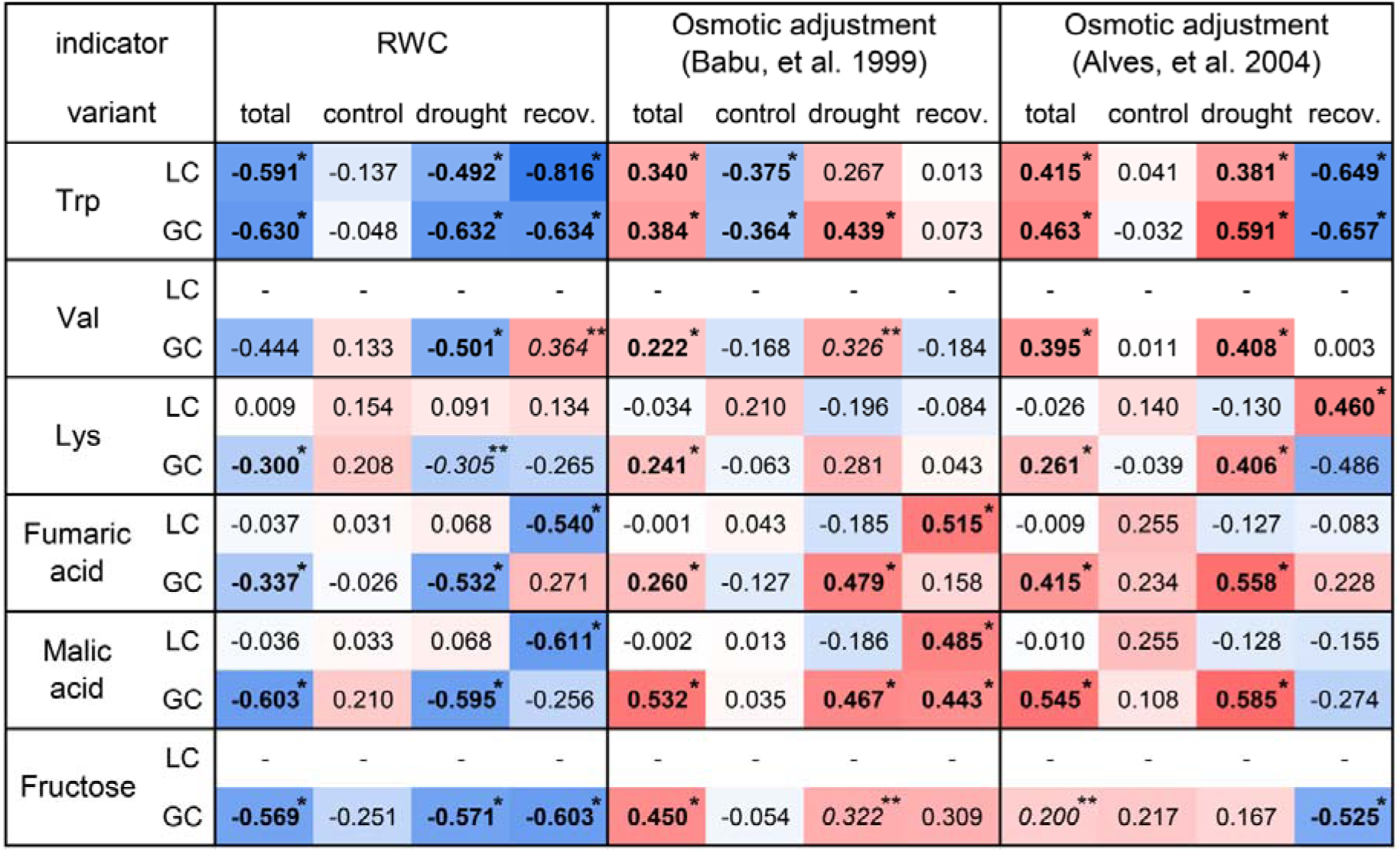
List of compounds demonstrating significant negative correlations with changes in RWC and in osmotic adjustments for total dataset, control, drought-stressed and recovering plants, according to non-targeted metabolomics analysis (LCMS and GCMS). Asterisks (*) denote significance of the Pearson’s correlation at *p*≤ 0.05, double asterisks (**) denote significance of the Pearson’s correlation at *p* ≤ 0.1

In total, only six of identified metabolites - tryptophan, valine, lysine, fumaric acid, malic acid and fructose – demonstrated significant (*p* ≤ 0.05) correlations with needles water status. Accordingly, these compounds were selected for subsequent absolute quantification to assess their contribution to the observed dynamics of metabolic adjustment. In addition to these six metabolites, we selected four compounds previously reported to be abundant in pine needles – namely ascorbic acid, shikimic acid, glucose, and sucrose (Pukacka and Pukacki, 2000; Rivas-Ubach *et al*., 2018; Zlobin *et al*., 2024) – were also subjected to targeted quantitative metabolomics analysis. Furthermore, given the importance of inorganic ions in maintaining cellular osmotic potential, the contents of three major inorganic cations (K^+^, Ca^2+^, Mg^2+^) in pine needles were quantified.

### 3.3. Absolute quantification of polar metabolites and major inorganic ions

The ten selected low-molecular-weight metabolites were quantified in needles of drought-treated, rewatered and control pine plants. Following to the earlier established scheme (Chantseva *et al*., 2019), the external calibration strategy appeared to be more suitable for the highly abundant metabolites (sucrose, malic acid, shikimic acid), whereas the standard addition approach was more applicable for the low-abundant metabolites (valine, fructose, glucose, lysine, tryptophan, fumaric acid). For ascorbic acid, despite its high abundance in pine needles, calibration using the standard addition method yielded the most reliable results. The standard addition is known to be more precise and is usually recommended, as it allows the most accurate estimation of the low-abundant target metabolites with consideration the accompanying matrix effects (Frenich *et al*., 2009). This was the fact in our study as well, but for highly-abundant metabolites external standardization gave better results.

The analysis revealed substantial differences in the needle contents of individual analytes. Fumaric acid, valine, lysine and tryptophan were found in the amounts of tens to hundreds nmol/g DW (Figure 4). Valine, lysine and tryptophan demonstrated rather similar accumulation patterns in pine needles. The contents of these metabolites (averaged here and elsewhere between lower and upper needles) increased promptly by the end of the treatment period in drought-stressed plants relative to the corresponding mean values of the dataset, as indicated by Z-scores exceeding 1.96 for all three compounds in the corresponding data point (critical *p*-value 0.025). By the end of the two week long experimental period, the differences between the drought-treated and control plants were 3.6-fold for valine (494.5 vs. 135.4 nmol/g DW), 2.15-fold for lysine (440.1 vs. 204.7 nmol/g DW), and 3.90-fold for tryptophan (1019.3 vs. 261.2 nmol/g DW). Consistent with the correlation analysis performed on the data matrix derived from the untargeted metabolomics experiment (see above), the amounts of all three compounds correlated significantly with changes in needle water status. Across the entire dataset, the contents of each amino acid were negatively correlated with RWC and positively correlated with both OA1 and OA2, with tryptophan contents exhibiting the highest correlation with the of water status parameters (*r* = -0.736 with RWC, 0.439 with OA1, and 0.467 with OA2). Surprisingly, when considering specifically plant under continuous drought, the only significant correlation was observed between Trp amount and RWC (*r* = -0.535). In the needles of the recovering plants, Trp and Lys contents correlated with both RWC and OA2, whereas Val content correlated only with RWC. In summary, among the three amino acids quantified, Trp demonstrated the most prominent increase by the end of the water stress period and the highest degree of correlation with the plant water status. Given that all three compounds were present in low concentrations (generally below than µmol/g DW), their estimated contribution to total cellular osmotic potential was minimal, not exceeding 1 kPa.

**Figure 4.**
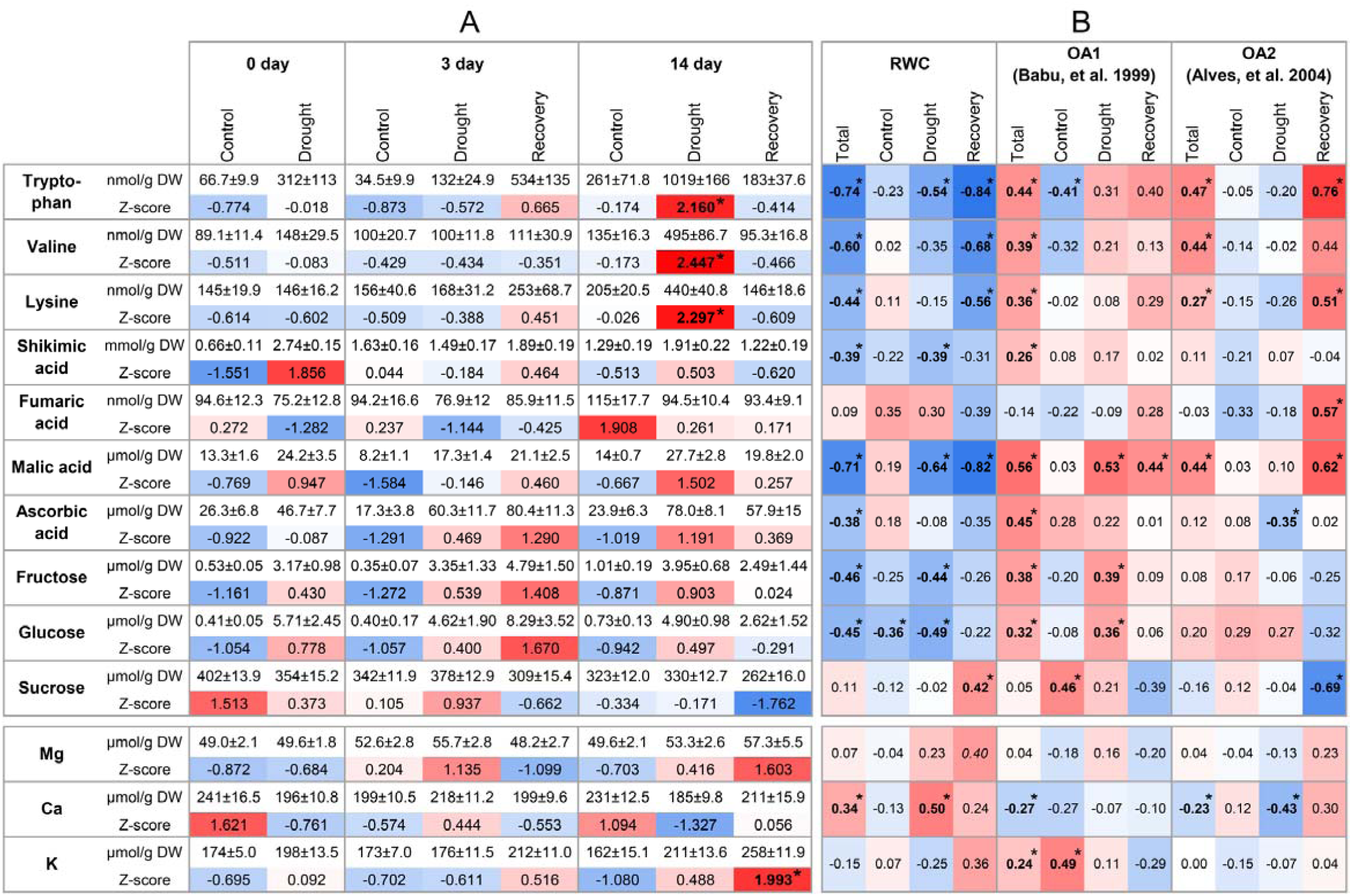
Absolute content (per g of DW), Z-scores (A) and correlations with RWC and osmotic adjustment 493 (OA1 and OA2) (B) for 10 quantified metabolites and for 3 inorganic ions. Contents of metabolites are given as arithmetic means, ± standard errors, Asterisks (*) denot significance of the Pearson’s correlation at *p* ≤ 0.05, or achievement of the z-score critical value at *p* ≤ 0.05

The accumulation patterns of fumaric acid was unrelated to water stress, with the most prominent accumulation observed in control plants by the end of the two-week period (Z-score 1.91) (Figure 4). No correlations of fumaric acid content with the parameters of needle water status were detected, with the sole exception of a correlation with OA2 in recovering plants (*r* = 0.568). As quantitative analysis demonstrated that fumaric acid levels were unresponsive to changes in water status, this compound was excluded from further analysis.

Fructose and glucose contents in needles ranged of several µmol/g DW. Both compounds demonstrated quite similar accumulation patterns, tending to accumulate in drought-stressed and recovering plants, although no significant Z-scores were observed for both metabolites (Figure 4). The highest contents of the both analytes were found in plants after three days of recovery, exceeding the contents in correspond control levels by 20.7-fold for glucose (8.29 vs. 0.40 µmol/g DW) and 13.7-fold for fructose (4.79 vs. 0.35 µmol/g DW). Both monosaccharides exhibited negative correlations with RWC and positive correlations with OA1 (but not OA2) across the entire dataset and when compared with the drought-stressed plants. For glucose, the negative correlation with RWC in control plants was also observed (*r* = -0.358). The combined tissue amounts of glucose and fructose were potentially sufficient to generate several kPa of osmotic pressure.

Ascorbic and malic acids were present in the needles at contents of tens µmol/g DW. Both acids tended to be more abundant in the needles of the drought-stressed and recovering plants compared to controls, although no significant Z-scores were observed for the both compounds (Figure 4). The highest ascorbic acid contents were found in the needles harvested on the third day of recovery, exceeding the control levels by 4.7-fold (80.4 vs. 17.3 µmol/g DW). In turn, the highest contents of malic acid was observed in drought-stressed plants by the end of the treatment, exceeding control plants 2.58-fold (27.7 vs. 14.0 µmol/g DW).

Malic acid contents exhibited clear correlations with the changes in the needle water status, evident in total dataset, in drought-stressed and in recovering plants. For ascorbic acid, the correlations with RWC and osmotic adjustment were less prominent, and even negative correlation with OA2 was observed in the drought-stressed plants (*r* = -0.353). The combined tissue concentrations of ascorbic and malic acids were potentially sufficient to generate several tens of kPa of osmotic pressure.

Finally, sucrose, shikimic acid and inorganic ions (K^+^, Ca^2+^, Mg^2+^) were present in the highest concentrations among the compounds studied, reaching tens and hundreds of µmol/g DW. However, among this group of major cell sap constituents, only shikimic acid demonstrated weak relations to the observed here changes in the needle water status, correlating slightly but significantly with RWC and OA1 in total data set and also with RWC in drought-stressed plants (Figure 4). For K^+^ and Mg^2+^, no consistent relationships with needle water status were observed, For sucrose and Ca^2+^, inverse relationships were detected in some instances, with positive correlations with RWC and negative correlations with OA (Figure 4).

## 4. Discussion

As confirmed by different calculation methods, the experimental drought treatment induced osmotic adjustment in pine saplings (Figure 1B). Also, by the end of the drought period (Day 14), the decline in RWC reached the levels associated with substantial dehydration of intracellular milieu (≈ 70%) (Figure 1A) (Lawlor and Cornic, 2002; Bartlett *et al*., 2012; John *et al*., 2018; Trueba *et al*., 2019). Thus, the experimental treatment was effective in eliciting both osmotic adjustment and significant cellular dehydration.

The combined application of LC-MS and GC-MS enabled the annotation of a total of 281 compounds in needle samples. However, only a limited subset of these metabolites exhibited drought-induced dynamics that correlated significantly with changes in needle water status. The analysis revealed similar accumulation patterns for three amino acids - valine, lysine, and tryptophan - which correlated with the dehydration-related changes of leaf water status (RWC and OA), and increased significantly by the end of the water stress period (Day 14). Given that a substantial drop in RWC was observed by this time point, it is plausible that these amino acids function as osmoprotectants in pine needles, specifically activated when the intracellular milieu becomes severely dehydrated. Accumulation of several amino acids under severe drought stress has also been reported in *Pinus radiata* (De Diego *et al*., 2013, 2015). Tryptophan is of particular interest, as it exhibited the most pronounced increase by the end of the drought period and the strongest correlations with both RWC and OA levels. Previous studies have shown that acclimation to freezing stress in conifers involves tryptophan accumulation, and that tryptophan may play a dominant role in the freezing stress response in conifers,- a pattern distinct from Arabidopsis, which relies primarily on proline accumulation (Angelcheva *et al*., 2014; Cañas *et al*., 2016). Our results indicate that tryptophan may also play a prominent role in the drought response, which is consistent with the fact that both severe drought and freezing stress lead to dehydration of the intracellular milieu (Beck *et al*., 2007). This observation aligns with a previous study on drought-treated Scots pine, which identified tryptophan as one of the most strongly up-regulated metabolites under drought conditions (MacAllister *et al*., 2019). It should be noted that drought-induced accumulation of tryptophan and valine has also been observed in angiosperms (Anzano *et al*., 2022). However, to the best of our knowledge, this study demonstrates for the first time in conifers a correlation between stress-induced dynamics of tryptophan content and plant water status.

Both glucose and fructose correlated with the changes in needle water status (RWC and OA), and tended not only to increase their tissue contents under water stress but also to maintain elevated levels during post-stress recovery. Drought-induced accumulation of sugars contributes to the protection of macromolecules under stress conditions. However, relatively unreactive non-reducing sugars are typically utilized as stress protectors, rather than monosaccharides such as glucose and fructose (Zulfiqar *et al*., 2020). The observed correlations with changes in water status may potentially indicate a role for glucose and fructose in osmotic adjustment, particularly in maintaining osmotic potential after stress relief. Nevertheless, their absolute quantities were relatively low and could account for no more than several kPa of osmotic pressure (Alves and Setter, 2004). Therefore, although the tissue contents of these monosaccharides followed changed in needle water status, their significance as osmolytes or osmoprotectants remains questionable.

In contrast to glucose and fructose, sucrose neither correlated with plant water status nor accumulated in drought-exposed plants. Indeed, positive correlations with RWC and negative correlations with OA were even observed for sucrose. This finding was unexpected, as sucrose is an abundant non-reducing sugar that can serve as an osmoprotectants (Sharma *et al*., 2021) and accumulate in cytosol as a compatible osmolyte. Accumulation of low-molecular-weight non-structural carbohydrates (mainly represented by sucrose) is responsible for routine osmotic adjustment at daily timescale (Rowland *et al*., 2023), and is frequently observed in water-stressed plants (De Diego *et al*., 2013; Hajek *et al*., 2022; Blumstein *et al*., 2023; Wang and Wang, 2023), as well as in plants during cold acclimation (Angelcheva *et al*., 2014; Chang *et al*., 2021), given that both drought and freezing stress negatively affect cellular hydration. The reasons for the absence of a correlation between sucrose accumulation and water deficit in the present study remain unclear.

Organic acids - ascorbic, malic and shikimic - were highly abundant in pine needles. Ascorbic acid content correlated with changes in water status, although the correlation was relatively weak. In contrast, malic acid exhibited correlations with RWC and OA changes in both drought-stressed and in recovering plants. Its relatively high abundance and strong correlation with water status suggest that malic acid may indeed play a role in osmotic adjustment processes in water-stressed pine plants. Shikimic acid demonstrated significant, albeit weak, correlation with changes in needle water status. However, given its high abundance - shikimic acid was the most abundant metabolite identified - even weak correlation with changes in cellular water status may indicate a meaningful contribution to the maintenance of osmotic potential. Inorganic cations (K^+^, Mg^2+^, Ca^2+^) did not exhibit any significant correlations with changes in water status, although accumulation of inorganic ions in a vacuole is a metabolically “cheap” strategy for lowering cellular osmotic potential without the need for organic solute synthesis.

In conclusion, the concurrent analysis of the drought-induced changes in the tissue water status and stress-dependent dynamics of individual polar metabolites enabled the identification of compounds likely involved in adaptive response to drought stress in Scots pine saplings. Tryptophan, and to a lesser extent valine and lysine, are likely involved in protection of cellular compounds during severe water stress, when RWC level drops below 70-75%. Accordingly, tryptophan biosynthesis may represent a potential target for enhancing ree drought tolerance in regions where severe droughts are frequent. Osmotic adjustment processes likely depend on the accumulation of organic acids, with malic acid demonstrating the clearest link to osmotic adjustment, and shikimic acid probably also playing an important role. Given that osmotic adjustment significantly contributes to the maintenance of both photosynthesis and growth under water-limited conditions (Zivcak *et al*., 2016; Blum, 2017; Turner, 2018), breeding strategies aimed at enhancing the water-responsive biosynthesis of these compounds hold promise. The original approach employed here – identifying key "soft" traits, such as the specific osmolytes and osmoprotectants characterized in this work – has substantial applied value, transforming drought tolerance from a vague concept into a set of quantifiable biochemical markers. This enables breeders to transition from slow, phenotype-based selection to marker-assisted selection targeting the specific metabolic pathways responsible for cellular protection and osmotic adjustment.

## Abbreviations

AAS: Atomic Absorption Spectrometry
AMDIS: Automated Mass Spectral Deconvolution and Identification System
DW: Dry Weight
FC: Fold Change
FDR: False Discovery Rate
GC-EI-Q-MS: Gas Chromatography-Electron Ionization-Quadrupole-Mass Spectrometry
GC-MS: Gas Chromatography-Mass Spectrometry
HPLC: High-Performance Liquid Chromatography
KNN: k-Nearest Neighbors algorithm
LC-MS: Liquid Chromatography-Mass Spectrometry
LC-MS/MS: Liquid Chromatography with Tandem Mass Spectrometry
LDR: Linear Dynamic Range
MRM: Multiple Reaction Monitoring
OA: Osmotic Adjustment
OA1: Osmotic Adjustment calculated by Method 1 (RWC-based)
OA2: Osmotic Adjustment calculated by Method 2 (Absolute WC-based)
Ψπ: Osmotic Potential
PCA: Principal Component Analysis
QqQ-MS/MS: Triple Quadrupole Tandem Mass Spectrometry
RI: Retention Index
RP-IP-HPLC: Reversed-Phase Ion-Pair High-Performance Liquid Chromatography
RWC: Relative Water Content
tR: Retention Time
WC: Water Content

## Acknowledgements

The metabolomic part of this work, including optimization of metabolite extraction, GC-MS and LC-MS analyses, and statistical analysis of the metabolome data, was supported by the Russian Science Foundation (Grant No. 25-14-00234). The plant material, experimental conditions, and analysis of mineral nutrient contents were supported by the Ministry of Science and Higher Education of the Russian Federation (Project No. 126012615950-2).

## Competing interests

The authors declare that they have no known competing financial interests or personal relationships that could have appeared to influence the work reported in this paper.

## Author contributions

AVK, IEZ, YVI: **conceptualization**; AVK, IEZ, YVI, AO, AS, TB, AF: **methodology**; AVK, IEZ, YVI, NF: **formal analysis**; AVK, IEZ, YVI, AII, AO, SS,: **investigation**; VVK, AF: **resources**; AVK, IEZ, YVI, AO, AS, TB: **data curation**; AVK, IEZ: **writing - original draft**; AVK, IEZ, YVI, AF: **writing - review & editing**; AVK and NF: **visualization**; VVK: **supervision and funding acquisition**

## Data availability

The data supporting the findings of this study are available from the corresponding author (A.V. Kartashov) upon request.

Appendix A. Supplementary material

## Notes

### Competing Interest Statement

The authors have declared no competing interest.

